# AVOCODO: An open-source multimodal annotation platform for developmental EEG

**DOI:** 10.64898/2026.08.27.747371

**Authors:** Winko W. An

## Abstract

Behavioral annotation of synchronized video recordings is an essential step in developmental electroencephalography (EEG) research, supporting both the identification of behavior-related artifacts and the investigation of brain–behavior relationships. Existing annotation workflows, however, are often fragmented: proprietary EEG software provides limited flexibility for behavioral coding, whereas dedicated behavioral annotation platforms typically lack native integration with EEG data. We developed AVOCODO (Audio/VideO CODing Optimization), an open-source MATLAB-based software platform that integrates synchronized behavioral annotation directly into the EEG workflow. AVOCODO reads native EGI MFF recordings, synchronizes embedded video with EEG, visualizes the audio spectrogram to facilitate precise annotation of vocalizations, and writes user-defined behavioral events directly back into the original MFF recording as native EEG event markers while simultaneously exporting annotations as CSV files. The software supports fully customizable behavioral coding schemes, optional EEG visualization for quality control, and reloading of previously annotated recordings for review and inter-rater verification. Since its initial development in 2024, AVOCODO has been applied internally across five developmental EEG studies involving approximately 500 pediatric participants and more than 3,000 EEG recordings. By bridging behavioral annotation and EEG preprocessing within a unified open-source workflow, AVOCODO has the potential to improve the efficiency, reproducibility, and scalability of behavioral annotation in developmental EEG research.

## 1. Introduction

Electroencephalography (EEG) has become an increasingly important tool in developmental neuroscience due to its portability, relatively natural recording environment, and suitability for studying populations that are difficult to assess using other neuroimaging modalities, such as infants, toddlers, and children with neurodevelopmental disorders (An et al., 2022, 2025; Barry-Anwar et al., 2024; Buzzell et al., 2023; Chung et al., 2026; Goodspeed et al., 2023).

Unlike functional magnetic resonance imaging (fMRI), which requires participants to remain motionless in a confined scanner, EEG recordings can be acquired while participants sit comfortably, often alongside their caregivers, and tolerate modest natural movement during data collection. Advances in EEG sensor technology have further improved participant comfort by reducing setup time and increasing the tolerability of prolonged recordings. Among commercially available systems, the EGI Geodesic Sensor Net has become one of the most widely adopted platforms in developmental EEG research owing to its rapid application, infant-friendly saline-based electrode design, availability of age-specific sensor nets, and support for high-density recordings. These features make it particularly well suited for studies involving infants, young children, and clinical populations.

Behavioral annotation is a critical component of the post-acquisition workflow in developmental EEG research. The EGI NetStation software, for example, supports synchronized video recording alongside EEG acquisition, enabling researchers to review participant behavior during offline analysis. These synchronized recordings serve at least two important purposes. First, participant behaviors during data collection can substantially affect EEG signal quality.

Behaviors such as talking, crying, pulling on the sensor net, or making large body movements introduce artifacts, while periods of inattention (e.g., looking away during a visual paradigm) reduce the signal-to-noise ratio and can compromise downstream analyses such as event-related potentials (ERPs). Second, many modern developmental EEG paradigms aim to investigate the relationship between neural activity and ongoing behavior. For example, studies of parent–child interactions often examine how neural activity in caregivers and children synchronizes during specific social behaviors (Turk et al., 2022). In such studies, accurately annotating behavioral events and aligning them with the EEG recording is essential for both artifact identification and behavioral-neural analyses.

However, the tools currently available for behavioral annotation each have important limitations. The EGI NetStation software provides an integrated environment for reviewing synchronized EEG and video recordings and allows users to manually insert behavioral markers directly into the EEG recording. While this workflow is familiar to many developmental EEG laboratories using EGI systems, it relies on proprietary software and a largely manual annotation process.

The interface is designed primarily for EEG review rather than efficient behavioral coding, making large-scale annotation time-consuming and offering limited flexibility for customized coding schemes or integration with modern open-source analysis pipelines. Consequently, some laboratories instead perform behavioral coding using dedicated annotation software, such as Datavyu (Datavyu: Video Coding and Data Visualization Tool, n.d.), Mangold INTERACT (Mangold INTERACT, n.d.), BORIS (Friard & Gamba, 2016), or ELAN (Sloetjes & Wittenburg, 2008), which provide richer interfaces for video annotation and support multiple annotation streams. However, these platforms are generally developed as standalone behavioral coding tools and have little or no native support for EEG data. As a result, behavioral annotations are typically exported as external files and subsequently synchronized with EEG recordings through custom scripts or manual processing steps. This fragmented workflow increases the complexity of data management, introduces additional opportunities for synchronization errors, and reduces the reproducibility and interoperability of behavioral annotation within modern EEG analysis pipelines.

To address these limitations, we developed AVOCODO (Audio/VideO CODing Optimization), an open-source MATLAB-based software platform for synchronized behavioral annotation of EEG recordings. AVOCODO is designed to combine the seamless EEG integration of the EGI NetStation workflow with the flexibility and efficiency of dedicated behavioral coding software. AVOCODO is built upon several well-established open-source software libraries. EEG data management is implemented using functions from EEGLAB (v2021.1) (Delorme & Makeig, 2004), while reading and writing of EGI MFF recordings is supported through MFFMatlabIO (v4.0) (Delorme, 2018/2026). Synchronized video playback is implemented using the open-source MATLAB–VLC interface developed by Léa Strobino (Strobino, 2019/2025), together with the VLC media player. The software provides a fully customizable annotation framework that allows users to define an arbitrary number of behavioral event types and tailor coding schemes to the needs of individual studies, rather than relying on predefined annotation categories. Upon completion of annotation, all user-defined behavioral events are written directly back into the original MFF recording as native EEG event markers, preserving compatibility with existing EGI analysis workflows. In addition, all annotations are exported as comma-separated values (CSV) files to facilitate downstream behavioral analyses and integration with modern open-source EEG processing pipelines.

## 2. Overview of AVOCODO

### 2.1 Software architecture and workflow

AVOCODO was developed in MATLAB and consists of two integrated workflows: an EEG processing workflow and a synchronized video/audio annotation workflow (Figure 1). The EEG workflow leverages EEGLAB (v2021.1) together with MFFMatlabIO (v4.0) to load, visualize, edit, and export EEG recordings in the EGI MFF format. The synchronized video/audio workflow integrates the MATLAB–VLC interface developed by Léa Strobino with the VLC media player to provide synchronized playback and behavioral annotation. During playback, users can annotate behavioral events based on visual observations, audio information, or both. These user-defined behavioral annotations are automatically incorporated into the original EEG recording as native event markers while preserving all existing events contained in the raw MFF file. In addition, AVOCODO can export all behavioral annotations as a CSV file for downstream behavioral analyses. To facilitate quality control, the software also provides optional visualization of the EEG time series and derived signal features, including power spectral density (PSD) and event-related potentials (ERPs), allowing users to rapidly assess data quality during the annotation process. Finally, AVOCODO supports reloading previously annotated recordings, enabling users to review, verify, and modify existing behavioral annotations without repeating the entire coding procedure.

**Figure 1.**
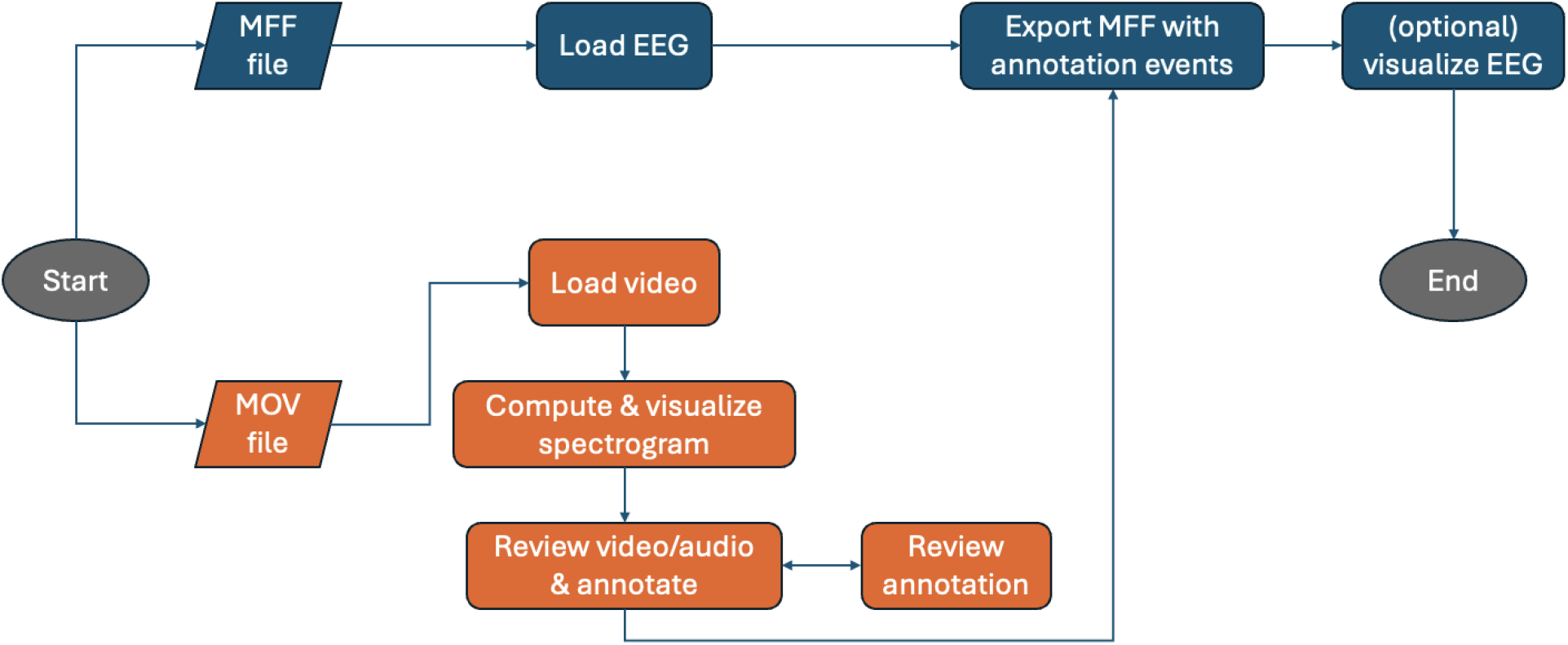
The AVOCODO workflow.

### 2.2 User interface

The AVOCODO user interface is shown in Figure 2. The interface is organized into several functional panels, including Load EEG and Video, Video Playback and Spectrogram, Coding, Markers, Export, and Plotting. To facilitate user onboarding, a toggle button located in the upper-left corner allows users to show or hide integrated instructions. When enabled, step-by-step instructions are displayed adjacent to the corresponding interface components. These instructions are sequentially numbered, allowing users to follow the recommended workflow in a straightforward manner (Figure 2).

**Figure 2.**
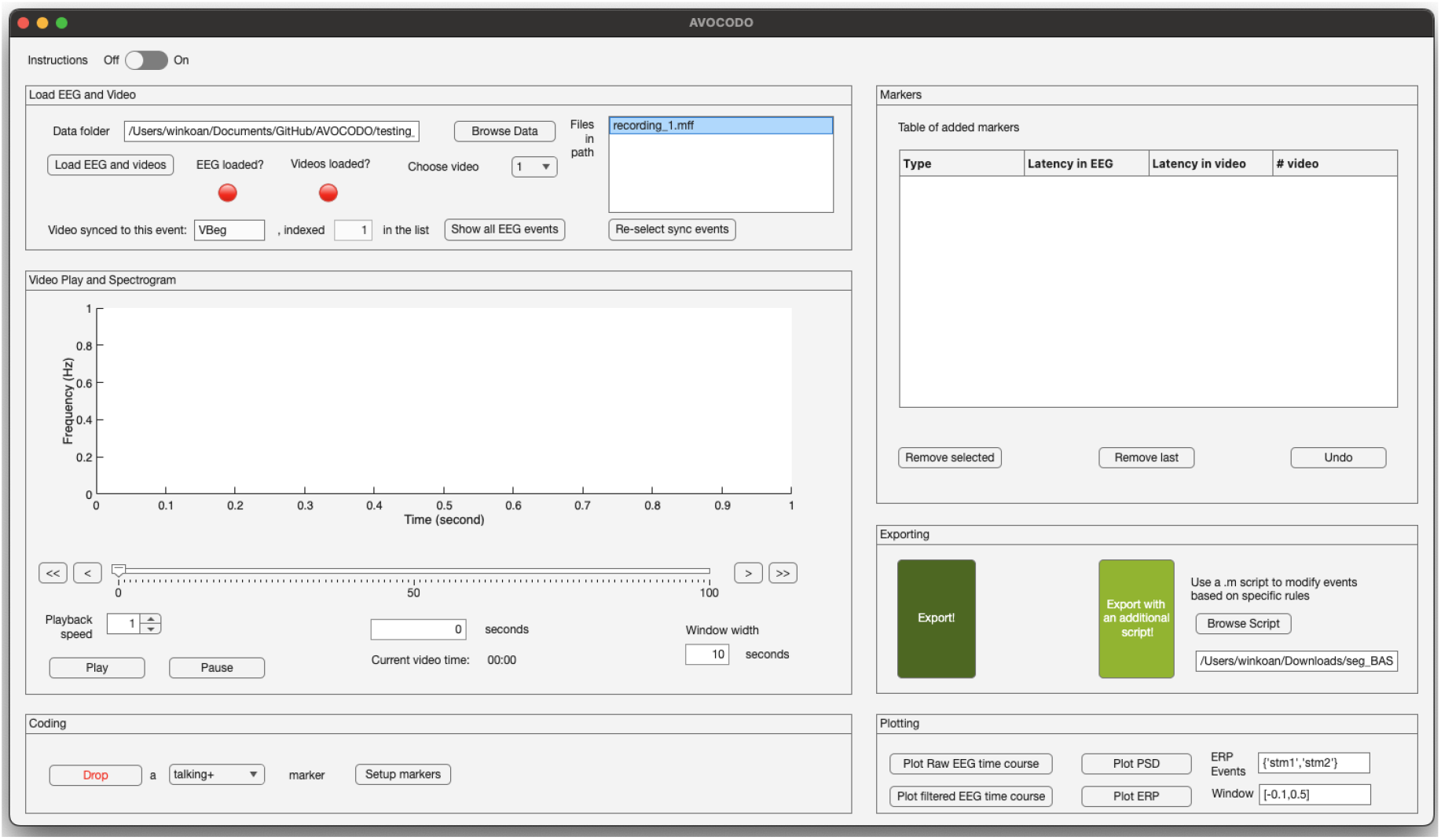
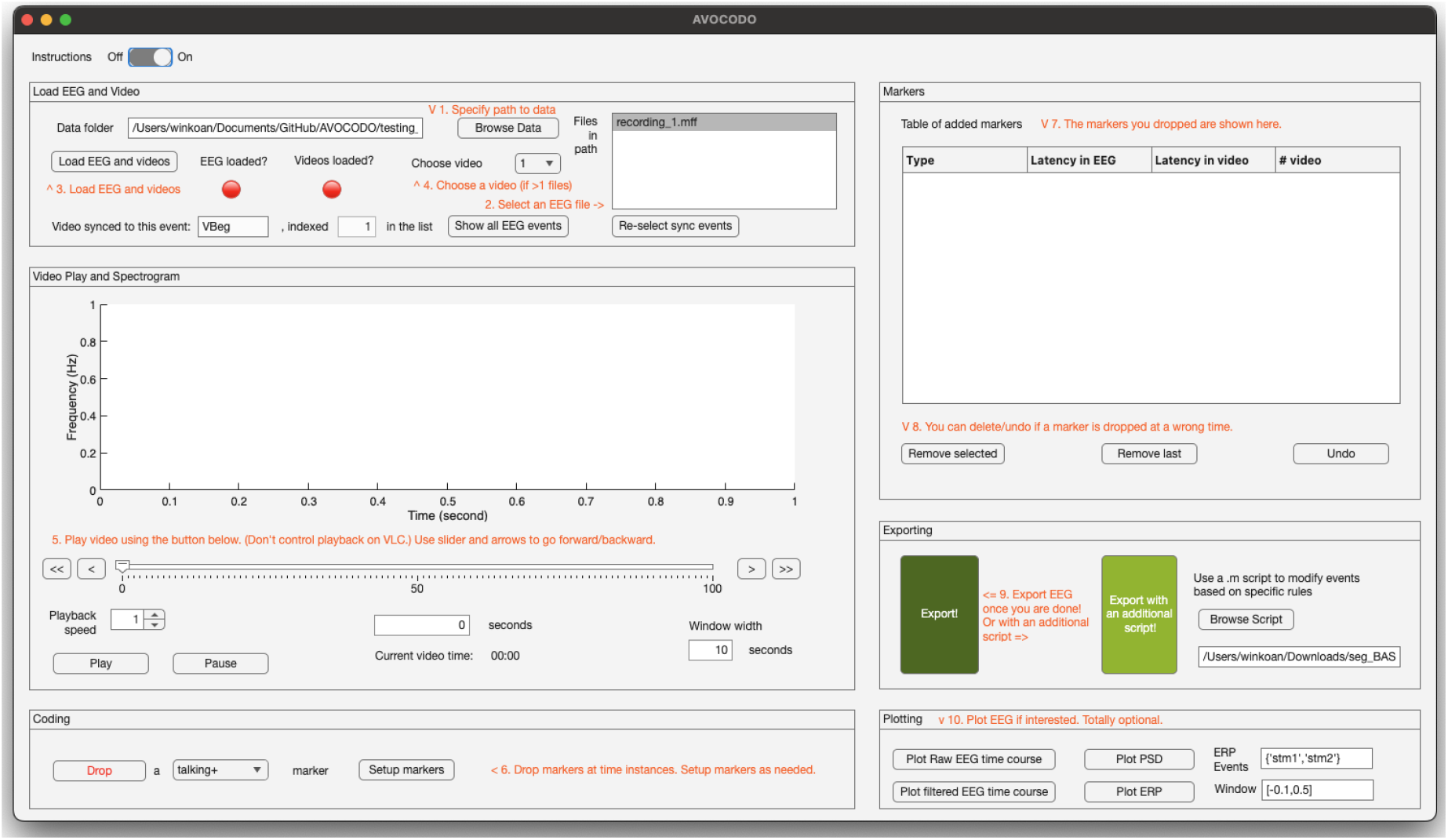
The AVOCODO user interface with instructions hidden (top) and displayed (bottom)

### 2.3 Data import

Users can specify the data directory either by entering the file path manually or by using the Browse Data button. Once a directory is selected, AVOCODO automatically searches for EGI MFF files within the folder. In the current release (v2.0), AVOCODO only supports MFF recordings acquired using the EGI NetStation software. Support for additional EEG file formats may be incorporated in future versions.

Because the embedded video and EEG streams have independent time bases, AVOCODO first identifies the synchronization events that link the two recordings. Upon successful loading, a dialog window prompts the user to select the existing event marker(s) that correspond to the onset of each synchronized video recording (Figure 3). These synchronization events are typically inserted automatically by NetStation at the beginning of each video recording and are used to align the EEG and video timelines. The number of synchronization events selected should match the number of video files embedded within the MFF recording. In the example shown in Figure 3, the MFF file contains two embedded MOV videos, and the user is therefore prompted to select two synchronization events. In this example, the event marker VBeg, which denotes the beginning of a video recording, is selected for both videos. Although VBeg is used here as an example, AVOCODO is compatible with any event marker name, provided that it correctly identifies the onset of the corresponding video recording. The software records the latency of the selected synchronization events and uses these timestamps to establish and maintain synchronization between the video and EEG throughout the annotation process.

**Figure 3.**
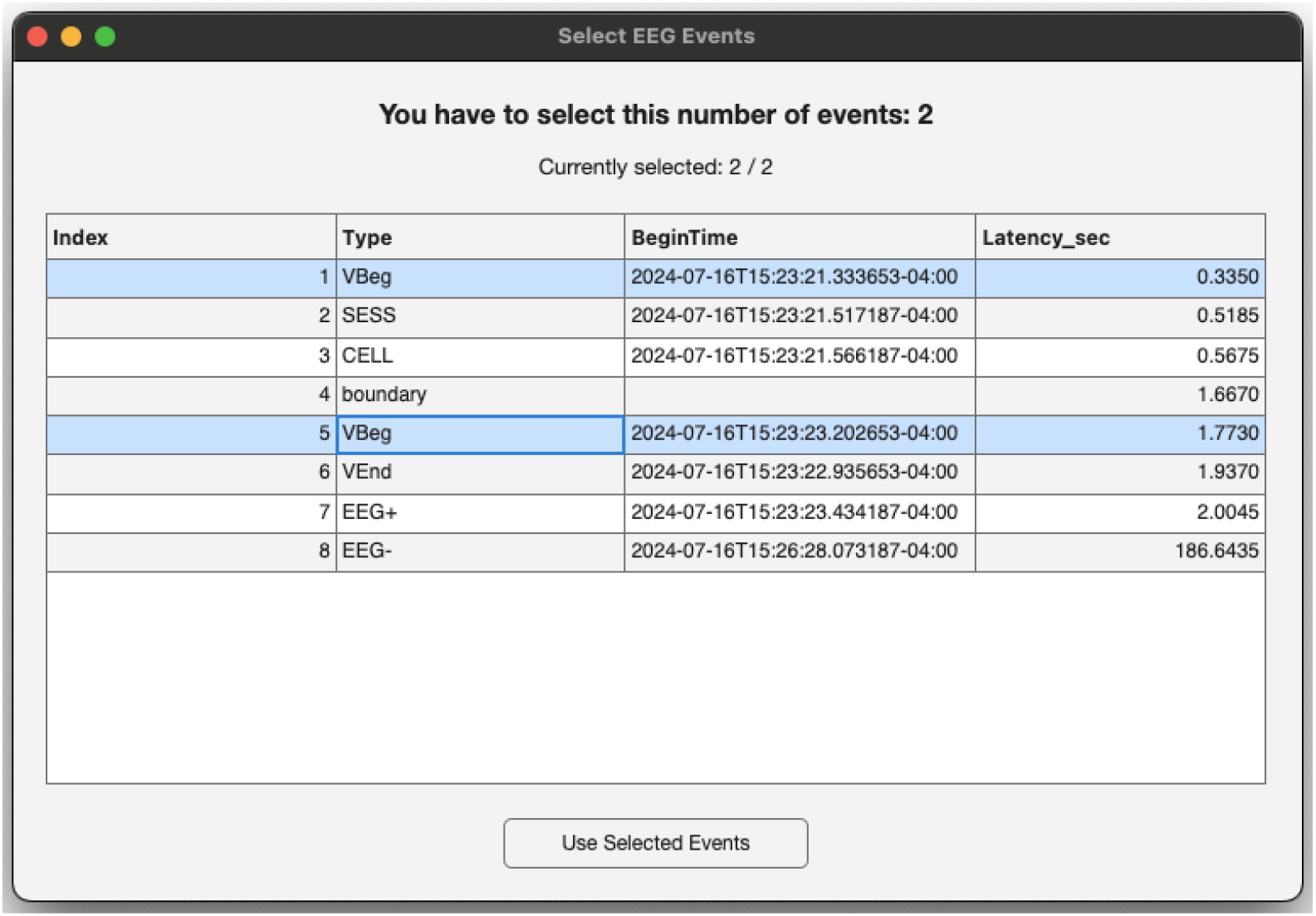
Window prompting the selection of EEG events for synchronizing with video(s)

### 2.4 Video Playback and Audio Spectrogram

Once both the EEG recording and synchronized video have been imported, AVOCODO displays the audio spectrogram of the selected video in the central panel, with existing EEG event markers overlaid according to their synchronized timestamps. The type and latency of each event are displayed adjacent to the corresponding marker for reference. Users can initiate synchronized video playback using the Play button within the Video Playback and Spectrogram panel. Clicking Play automatically launches the VLC media player while maintaining synchronization between the video, audio spectrogram, and EEG event timeline throughout playback. Users can control the current playback position, playback speed, and playback functions—including pause and forward/backward navigation—using the buttons within the Video Playback and Spectrogram panel.

**Figure 4.**
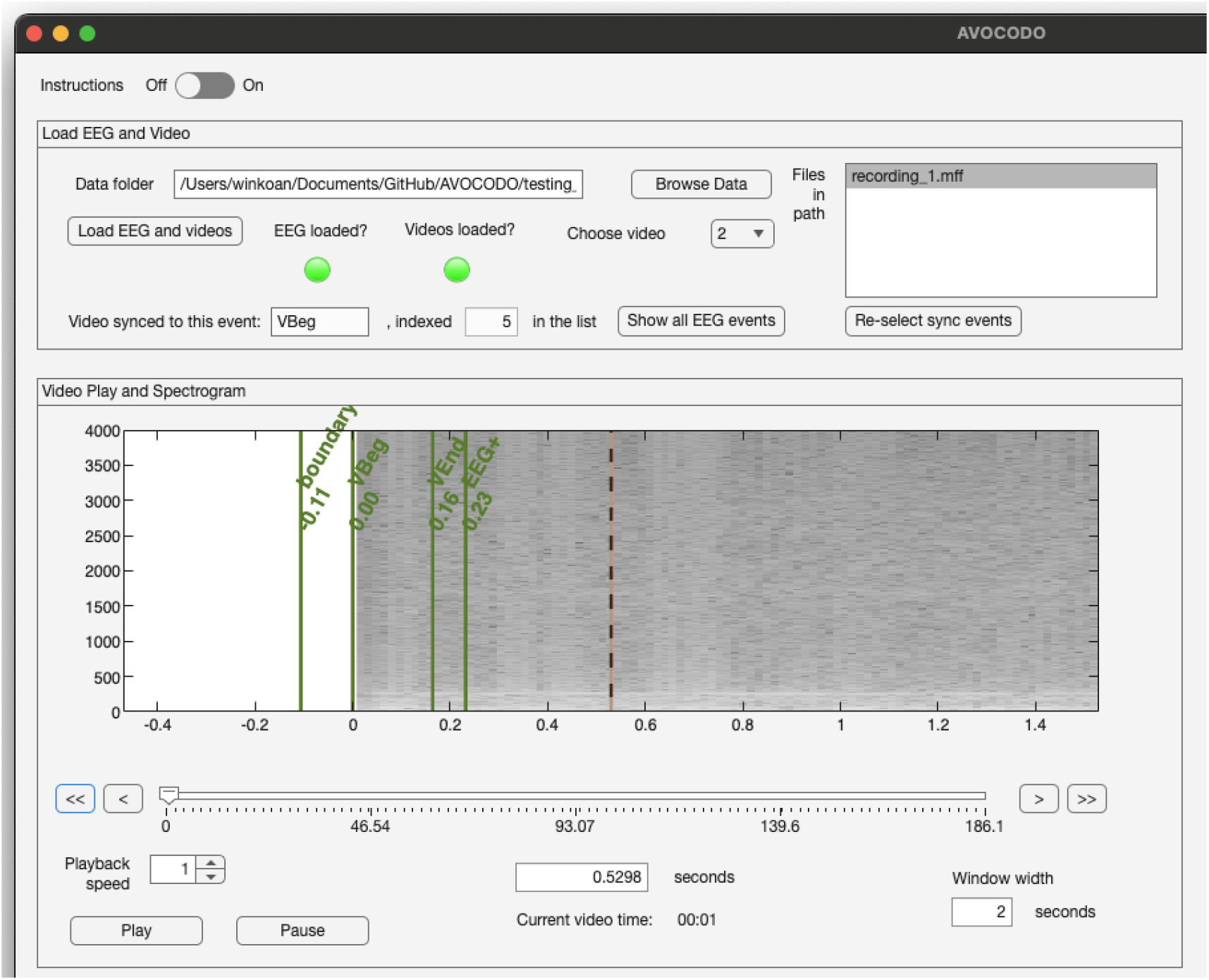
Once the EEG and video have been imported, the audio spectrogram is displayed in the central panel, with existing EEG event markers overlaid for reference.

**Figure 5.**
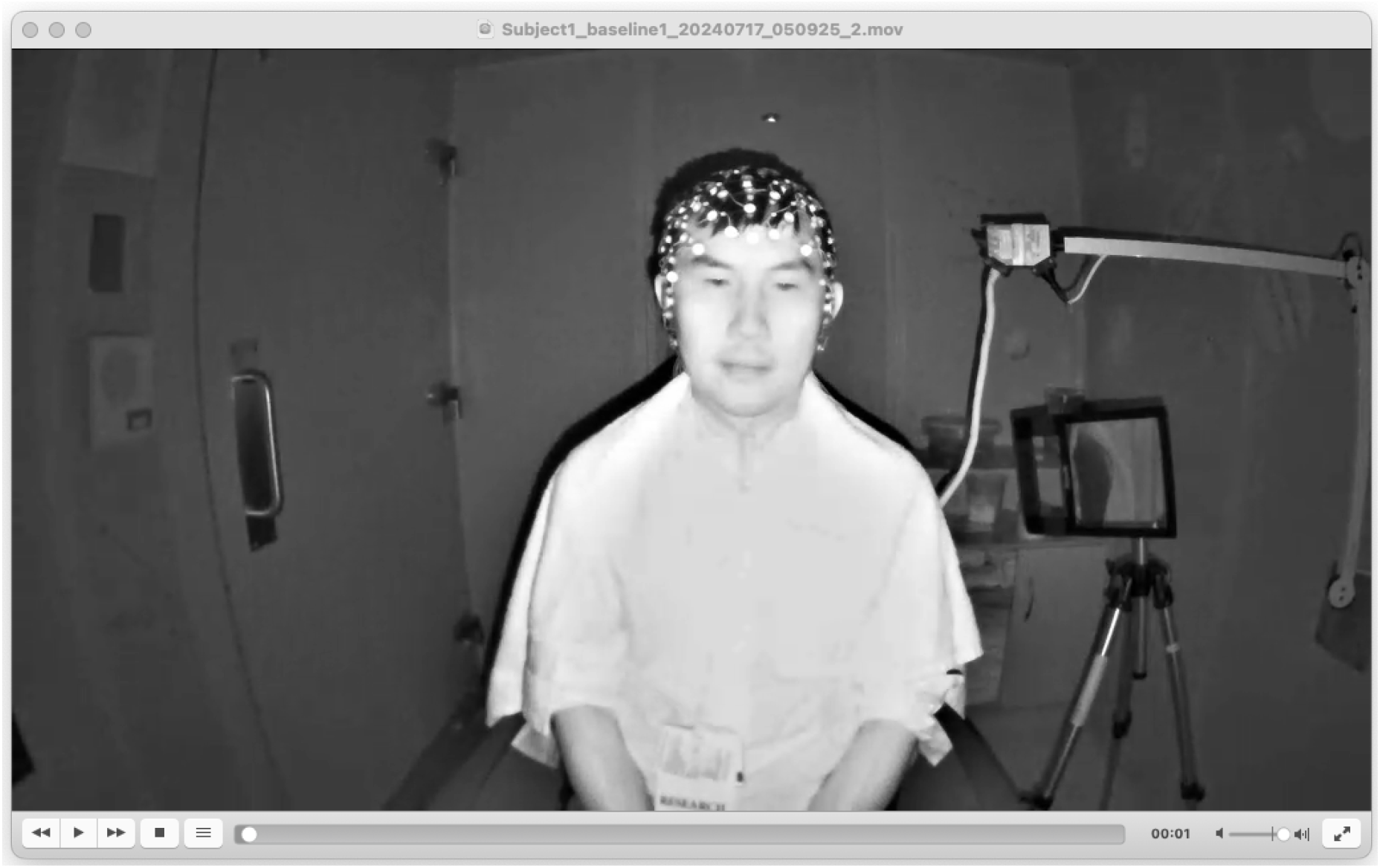
VLC is automatically launched once the Play button is clicked. The participant shown in the video is the author and has provided consent for the use of this image in the manuscript.

### 2.5 Flexible annotation framework

Before behavioral annotation begins, users define the event markers corresponding to the behavioral categories they wish to annotate. Clicking the Setup Markers button within the Coding panel opens a dialog window for configuring the behavioral annotation markers (Figure 6). Marker names are entered as comma-separated text strings. In the example shown, the onset and offset of each behavior are denoted by the suffixes “+” and “−”, respectively.

**Figure 6.**
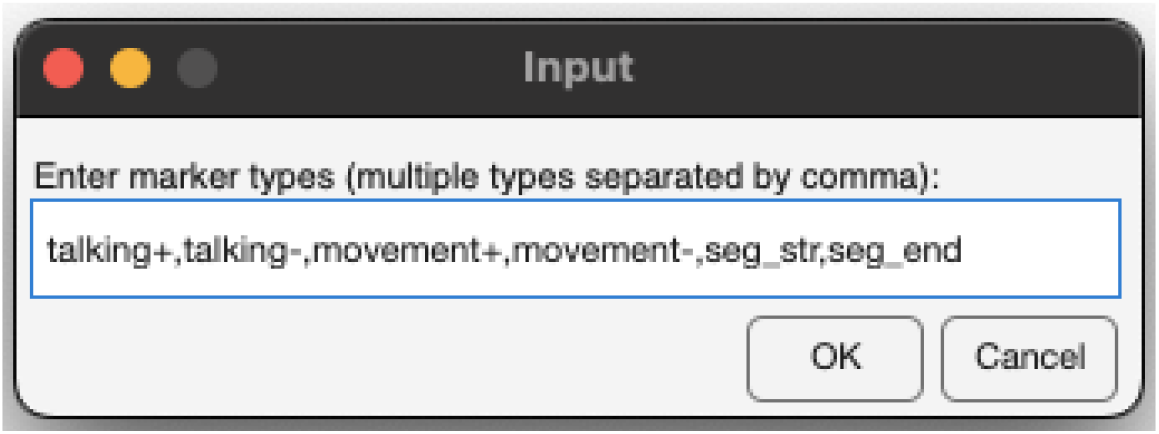
Dialog window for configuring behavioral annotation markers.

AVOCODO imposes no restrictions on the number or naming of markers, allowing users to define arbitrary behavioral categories and annotation schemes according to the requirements of their study.

### 2.6 Behavioral annotation workflow

Behavioral annotation in AVOCODO is performed using synchronized video and audio information. Behaviors such as sensor net pulling, head movements, and other visible participant actions can be identified from the video, while speech, crying, and other vocalizations can be identified from both the video and audio. In particular, the audio spectrogram displayed in the central panel facilitates precise annotation of speech and vocalization onset (Figure 7). To annotate an event, users select the desired marker from the drop-down menu in the Coding panel and click the Drop button to place the marker at the current playback position. The newly created annotation is immediately added to the event table in the Markers panel, together with its corresponding EEG latency, video timestamp, and the source video from which the annotation was created. Selecting an annotation from the table automatically navigates the video to the corresponding time point, enabling rapid review and verification. Incorrectly placed annotations can be removed or modified using the controls provided in the Markers panel.

**Figure 7.**
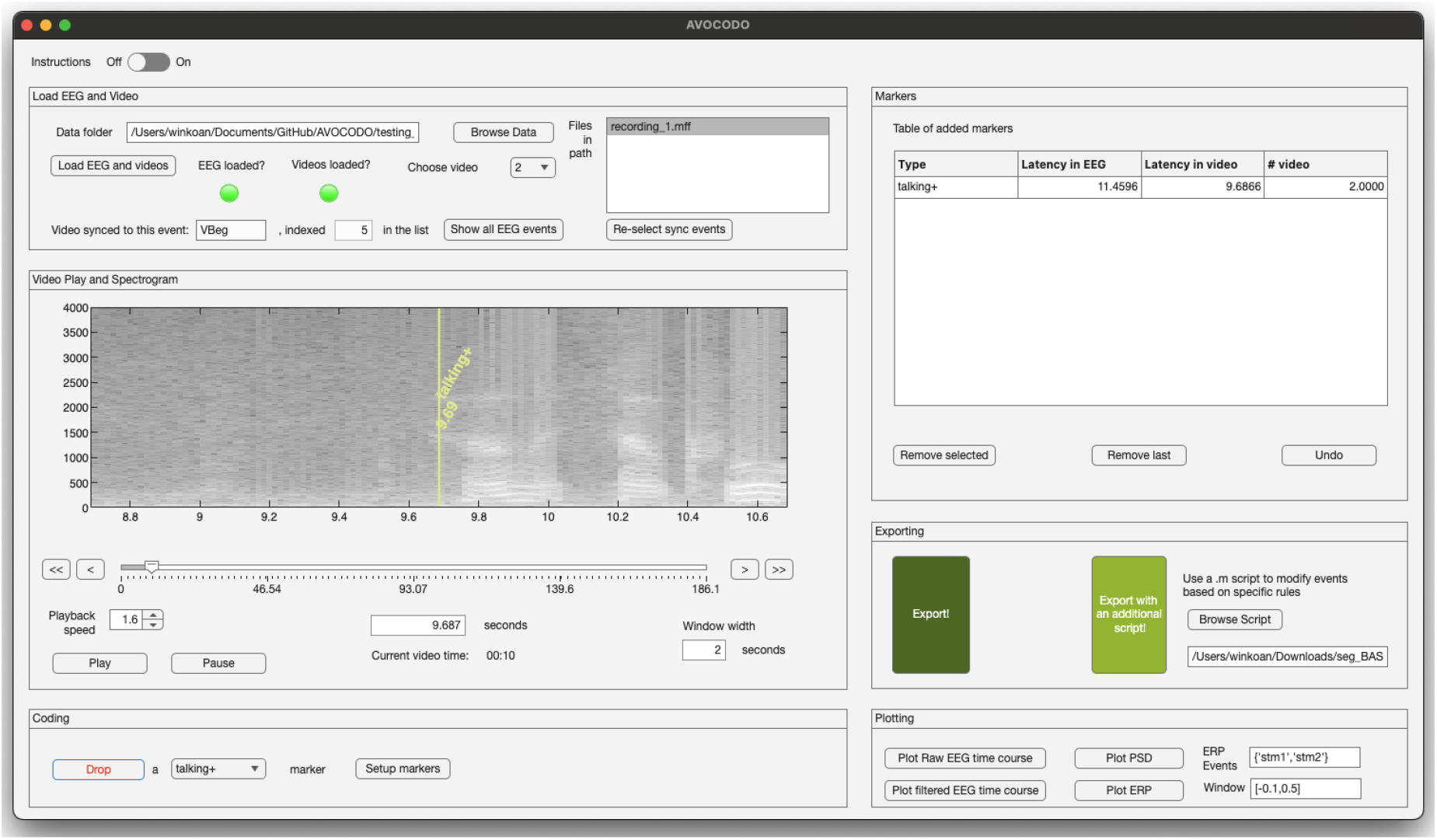
Precise annotation of speech/cry onset using the audio spectrogram.

### 2.7 Data export

Once behavioral annotation is complete, users can export the results by clicking the Export button within the Export panel. AVOCODO simultaneously generates two outputs: (1) an annotated MFF file containing all behavioral annotations embedded as native EEG event markers, and (2) a CSV file containing the behavioral annotation data for downstream behavioral analyses. To facilitate data organization, the software automatically creates two subdirectories within the selected data folder: 0_markers, which stores the annotation CSV files, and 1_marked_EEG, which stores the annotated MFF recordings.

For advanced users, AVOCODO also provides an optional custom export script that is executed immediately before the annotated EEG is written to the MFF file. This feature allows users to perform study-specific modifications to the EEGLAB “EEG” structure, including the event information stored in “EEG.event”, prior to export. For example, users may choose to remove existing EEG event markers that occur within annotated behavioral intervals, relabel events, or perform other customized processing steps before saving the final MFF file. Because these operations are often highly specific to individual experimental protocols, AVOCODO intentionally provides this functionality as a user-editable MATLAB script, enabling researchers to tailor the export procedure to the needs of their own studies while preserving the core annotation workflow.

### 2.8 Closing the interface

When the interface is closed normally using the close button in the main window, AVOCODO automatically saves the current session configuration to the “default.mat” file located in the “config” directory. The saved configuration includes user-defined settings such as the data directory, behavioral marker definitions, and any custom export script. When AVOCODO is launched in a subsequent session, these settings are automatically restored, allowing users to resume annotation without repeating the configuration process and thereby improving workflow efficiency.

### 2.9 Data visualization

A key design principle of AVOCODO is the separation of behavioral annotation from EEG signal interpretation. Behavioral events are annotated solely on the basis of the synchronized video and audio recordings, ensuring that the annotation process is not influenced by the quality or characteristics of the EEG signals. This separation helps maintain the objectivity of behavioral coding while allowing EEG processing to be performed independently at a later stage.

Nevertheless, AVOCODO also provides optional EEG visualization tools to facilitate data quality assessment during the annotation process. At any point, users may inspect the EEG using the controls within the Plotting panel. The software supports visualization of the raw and bandpass-filtered EEG time series, power spectral density (PSD; Figure 8), and event-related potentials (ERPs), which are generated based on the event type and time window specified by the user. These visualization tools are intended solely to assist with quality control and are not required for behavioral annotation.

**Figure 8.**
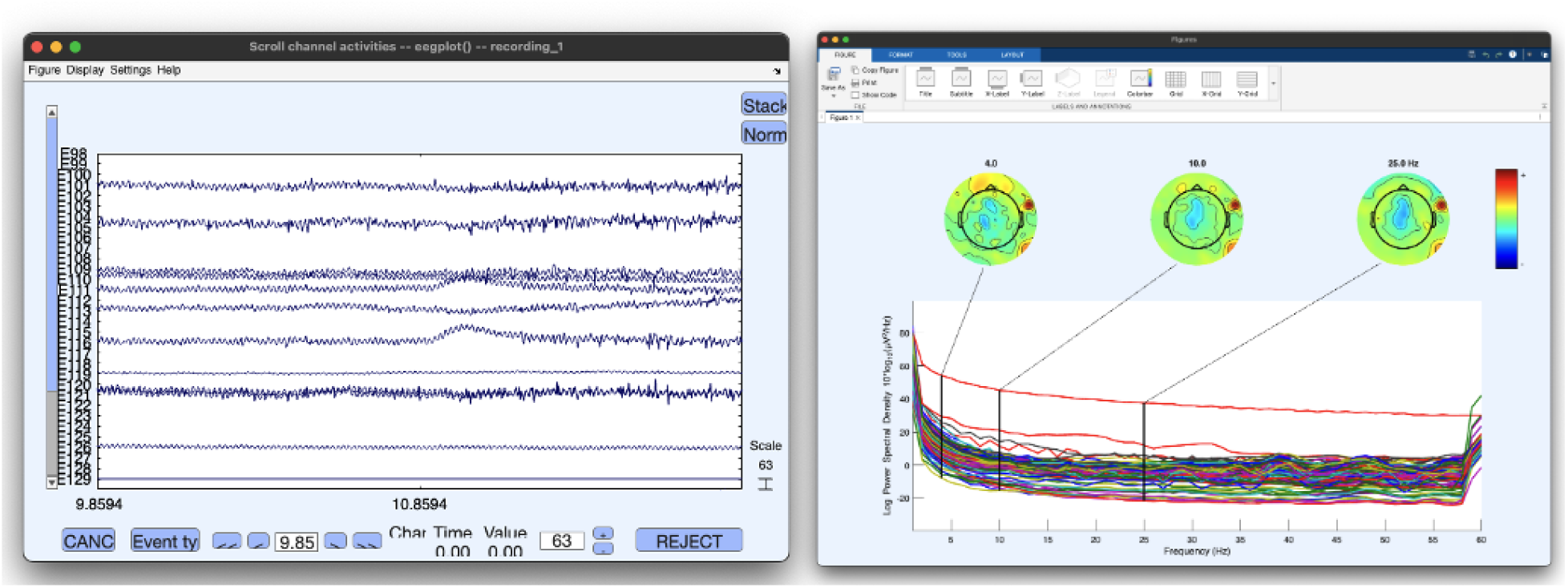
Optional EEG visualization tools provided by AVOCODO. Left: EEG time-series visualization. Right: Power spectral density (PSD) visualization.

### 2.10 Integration into preprocessing pipeline

An important consideration for an EEG-centered behavioral annotation tool is its integration with downstream EEG preprocessing and analysis workflows. Rather than exporting behavioral annotations as external files that require subsequent synchronization, AVOCODO writes all user-defined behavioral events directly into the original MFF recording as native EEG event markers. By preserving the original MFF file format, annotated recordings can be incorporated seamlessly into existing preprocessing pipelines without requiring additional file conversion or synchronization steps. This design allows behavioral annotations to become an integral part of the EEG recording while maintaining full compatibility with software and analysis pipelines that support the MFF format.

### 2.11 Loading an annotated file

Another important consideration for behavioral annotation is the ability to review and verify previously generated annotations. AVOCODO supports reloading of previously annotated EEG recordings, provided that the corresponding annotation CSV file is available in the output directory. Upon loading, all previously created behavioral annotations are restored, allowing a reviewer to inspect or modify the existing annotation set. Reviewers may either perform an independent annotation of the recording following the original coding protocol or navigate directly to previously annotated events by selecting entries in the Markers table. Selecting an event automatically synchronizes the EEG and video to the corresponding time point, enabling efficient verification, correction, or removal of individual annotations as needed.

## 3. System Requirements and Software Availability

AVOCODO has been tested with MATLAB releases R2019a through R2025a and VLC media player version 3.0.16 through 3.0.24 on both Windows and macOS platforms. The software incorporates EEGLAB (v2021.1) for EEG data management, MFFMatlabIO (v4.0) for reading and writing EGI MFF files, and the open-source MATLAB–VLC interface developed by Léa Strobino for synchronized video playback. AVOCODO is distributed with these software components where permitted by their respective licenses, allowing installation without additional software configuration beyond MATLAB and VLC.

AVOCODO is released as open-source software under the Apache License 2.0 and is freely available on GitHub at https://github.com/winkoan/AVOCODO. The repository includes documentation, installation instructions, example datasets, and the source code.

## 4. Current applications of AVOCODO

Since its initial development in mid-2024, AVOCODO has been used internally for behavioral annotation in five developmental EEG studies involving approximately 500 pediatric participants and more than 3,000 EEG recordings across a diverse range of experimental paradigms. These studies demonstrate the software’s robustness across multiple experimental designs, with several manuscripts currently in preparation.

Figure 9 presents an example application of AVOCODO in a developmental EEG study examining visual evoked potentials (VEPs) in children with and without autism spectrum disorder. In the recording shown, a 5-year-old male participant with autism displayed approximately 5 s of crying and 81 s of visual aversion during a 3-min VEP experiment (Figure 9A). These behaviors were annotated using AVOCODO and subsequently used to exclude the corresponding EEG segments during preprocessing with the HAPPE automated preprocessing pipeline (Gabard-Durnam et al., 2018; Lopez et al., 2022). Without incorporating behavioral annotations, preprocessing retained 282 VEP trials. After excluding EEG segments corresponding to behaviorally annotated intervals, 195 trials remained for analysis, indicating that 87 trials (30.9%) were associated with periods of crying and visual aversion.

**Figure 9.**
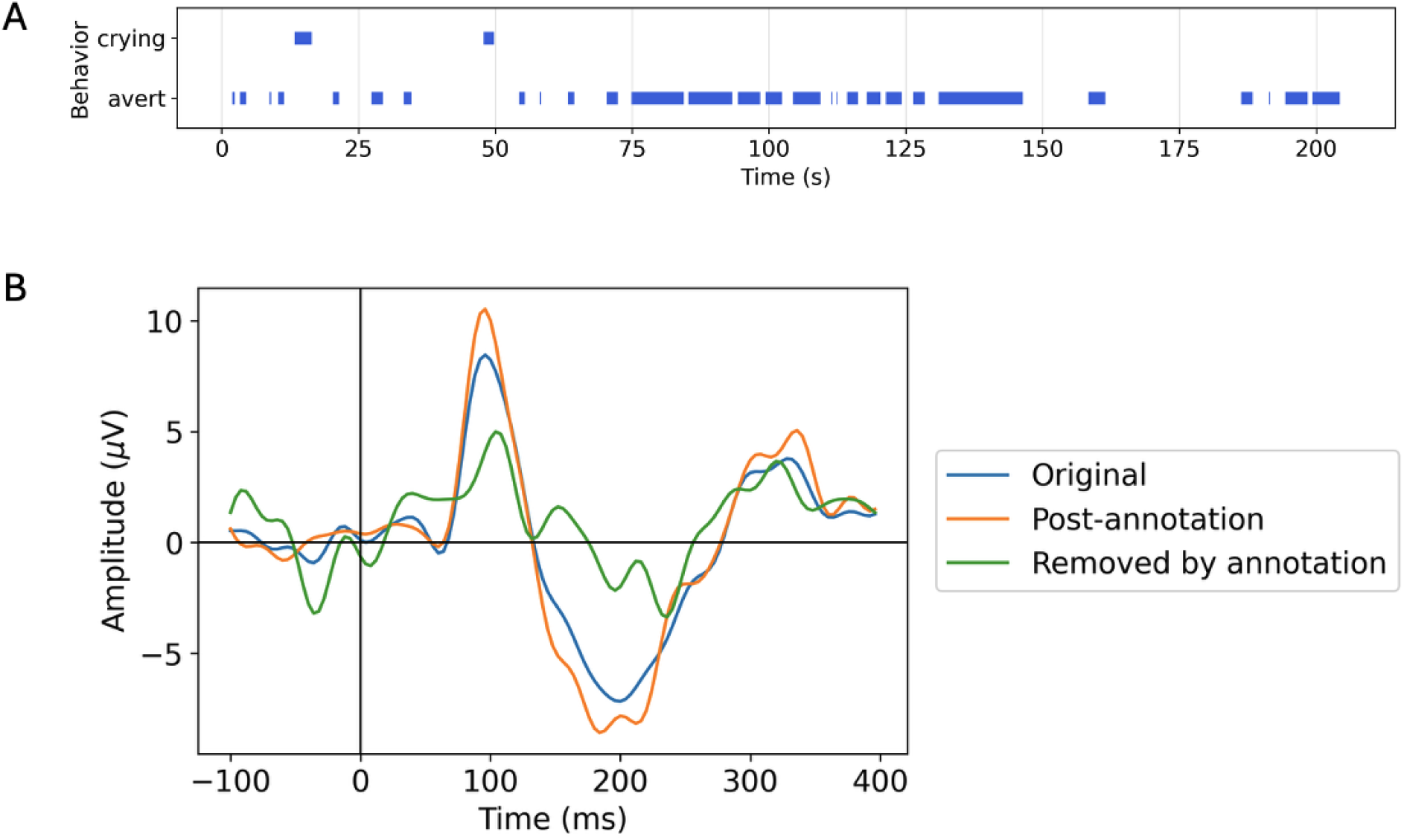
One example case of applying AVOCODO in VEP analysis

Figure 9B compares the average VEP waveforms derived from three sets of trials: (1) all retained trials without behavioral annotation (282 trials), (2) the retained trials after excluding behaviorally annotated segments (195 trials), and (3) the excluded trials alone (87 trials).

Compared with the waveform generated without behavioral annotation, the VEP obtained after excluding behaviorally annotated segments exhibits stronger P1 (∼100 ms) and N2 (∼200 ms) components. In contrast, the average waveform computed from the excluded trials differs markedly from the retained VEPs and lacks the characteristic morphology of a typical VEP, suggesting that these trials were substantially affected by behavioral or movement-related contamination. Although this example is intended to illustrate the practical application of AVOCODO rather than serve as a formal validation study, it demonstrates how synchronized behavioral annotation can facilitate the identification and exclusion of behaviorally contaminated data, thereby improving the quality and interpretability of downstream EEG analyses.

## 5. Contribution to research

Compared with representative software currently used for behavioral annotation in developmental EEG research (Table 1), AVOCODO combines the native EEG integration provided by EGI NetStation with the flexibility of dedicated behavioral annotation software. This integrated design addresses several methodological and practical limitations of existing workflows.

**Table 1.** Comparison of representative behavioral annotation tools commonly used in developmental EEG research. AVOCODO combines native EEG integration with the flexibility of dedicated behavioral annotation software while remaining open source.

| Workflow component | NetStation | Datavyu | BORIS | ELAN | INTERACT | AVOCODO |
| --- | --- | --- | --- | --- | --- | --- |
| Review synchronized EEG | ✓ | X | X | X | X | ✓ |
| Review synchronized video | ✓ | ✓ | ✓ | ✓ | ✓ | ✓ |
| Audio spectrogram | X | X | X | Limited | X | ✓ |
| Flexible behavioral coding | Limited | ✓ | ✓ | ✓ | ✓ | ✓ |
| Native EEG event generation | ✓ | X | X | X | X | ✓ |
| Export behavioral table | Limited | ✓ | ✓ | ✓ | ✓ | ✓ |
| Open source | X | ✓ | ✓ | ✓ | X | ✓ |

A major advantage of AVOCODO is its open-source design. Unlike proprietary software that requires institution-specific licenses and installation on designated laboratory computers, AVOCODO can be freely deployed on any computer with a MATLAB installation. Furthermore, the software can be compiled as a standalone executable using the MATLAB Runtime, allowing it to operate without a MATLAB license. This flexibility enables behavioral annotation to be distributed across multiple annotators, including students and research assistants, without being constrained by software licensing or access to dedicated workstations. Consequently, annotation can be performed more flexibly across laboratories and collaborative research groups.

AVOCODO is designed to integrate behavioral annotation directly into the EEG processing workflow. Whereas general-purpose behavioral annotation software typically exports behavioral events as external annotation files that must subsequently be synchronized with EEG recordings, AVOCODO directly reads native EGI MFF recordings and writes behavioral annotations back into the original EEG file as native event markers. As a result, behavioral annotations become an integral component of the EEG recording and are immediately compatible with downstream preprocessing and analysis pipelines that support the MFF format. At the same time, AVOCODO exports behavioral annotations as CSV files, allowing the same annotation data to be incorporated into behavioral analyses independent of the EEG workflow. This dual-output strategy eliminates redundant synchronization steps, reduces opportunities for human error, and promotes reproducible data processing.

AVOCODO enhances the temporal precision of behavioral annotation by combining synchronized EEG, video, and audio with visualization of the corresponding audio spectrogram. Precise identification of speech, crying, and other vocalizations can be challenging when relying solely on video playback, particularly when vocalization onset is subtle or partially obscured by background noise. The synchronized spectrogram provides a visual representation of the acoustic signal, enabling annotators to identify vocalization onset and offset with greater temporal accuracy. Improved behavioral timestamps facilitate more accurate alignment between behavioral events and neural activity, benefiting analyses including event-related potentials, time-frequency analyses, hyperscanning, and brain–behavior synchronization.

Behavioral annotation is inherently iterative and frequently involves multiple annotators, repeated quality-control procedures, and consensus review. AVOCODO supports this workflow by allowing previously annotated datasets to be reloaded, reviewed, modified, and independently reannotated while maintaining synchronization between the EEG recording, video, and behavioral events. These capabilities facilitate inter-rater reliability assessment, consensus coding, and transparent documentation of behavioral annotations, thereby improving the reproducibility of behavioral coding in developmental EEG research.

## 6. Limitations and future directions

Despite its advantages, AVOCODO has several limitations that provide opportunities for future development. First, the current version is designed specifically for EEG recordings acquired using the EGI NetStation system and the native MFF file format. Although MFF is widely used in developmental EEG research, investigators using other acquisition systems, such as BrainVision, BioSemi, or EDF-based workflows, are not currently supported. Extending AVOCODO to additional EEG file formats would broaden its applicability to a wider range of neuroscience laboratories while preserving the same integrated annotation workflow.

Second, AVOCODO is currently implemented as a MATLAB-based application. Although MATLAB is widely available in academic research institutions and standalone executables can be generated using the MATLAB Runtime, future versions could further improve accessibility through platform-independent implementations or web-based interfaces that facilitate collaborative annotation and deployment across multiple operating systems.

Finally, AVOCODO currently focuses on manual behavioral annotation. This design was chosen deliberately because behavioral coding in developmental neuroscience often requires expert judgment when interpreting complex behaviors, particularly in infants and children.

Nevertheless, the modular architecture of AVOCODO provides a foundation for future integration of automated behavioral coding algorithms. The current workflow can be readily extended by incorporating computer vision, speech recognition, or other machine learning approaches to generate preliminary behavioral annotations before manual review. Rather than replacing human annotators, such automated methods could provide initial event proposals that are subsequently verified and corrected by trained coders. This human-in-the-loop approach has the potential to substantially reduce annotation time while maintaining the accuracy and reliability required for developmental neuroscience research. As machine learning methods for behavioral analysis continue to mature, AVOCODO provides a flexible framework for integrating these technologies into existing EEG annotation workflows.

We envision AVOCODO as an evolving open-source platform that grows alongside advances in behavioral neuroscience and artificial intelligence. By combining native EEG integration with an extensible annotation framework, future versions have the potential to support increasingly efficient, reproducible, and scalable behavioral annotation for developmental EEG research.

## Acknowledgements

The author gratefully acknowledges colleagues who provided valuable feedback, tested early versions of the software, and offered suggestions throughout the development of AVOCODO, including, but not limited to, Virginia Rosenberger, Brooke Keough, Meagan Tsou, and Michael Khela.

